# BioTransformer 4.0 a comprehensive computational tool for small molecule metabolism prediction

**DOI:** 10.1101/2025.07.28.667289

**Authors:** Siyang Tian, Yannick Djoumbou Feunang, Eponine Oler, Fei Wang, Russell Greiner, Emma H. Palm, Emma L. Schymanski, Claudine Manach, David S. Wishart

## Abstract

BioTransformer 4.0, the successor to BioTransformer 3.0, is a freely available *in silico* metabolism prediction tool. It integrates both knowledge-based and machine learning approaches to predict metabolites for small molecules using one of seven modules: abiotic, environmental, CYP450, phase II, enzyme commission-based, human gut microbial, and all human metabolism. It also provides a customizable sequence prediction module that allows users to simulate multi-step metabolic transformations by chaining among the first six different modules.

BioTransformer 4.0 can make predictions more efficiently and accurately than the previous version, as it includes more than 130 new reaction rules, and also an optional validation module to improve the efficiency by restricting the number of predicted metabolites, due to their similarity among real human metabolites. We evaluated its performance by running the six-step all-human metabolism prediction on the DrugBank dataset of 2,457 known biotransformations, and the PhytoHub dataset of 633 known biotransformations – achieving recall values of 87.2% (resp., 91.6%) for the DrugBank (resp., PhytoHub) datasets.

## 1 Introduction

The exploration and identification of metabolites of small molecules plays a critical role in diverse fields such as drug discovery, environmental toxicology, metabolomics, and systems biology. Understanding how compounds are transformed within the human body and in microbial environments can guide early-stage pharmacokinetic profiling, toxicity assessments, and environmental pollutant analysis [9]. Traditional metabolite identification typically involves exploring plausible metabolite structures both theoretically and experimentally, which is time-consuming, expensive, and often incomplete [6].

To overcome these limitations, several *in silico* tools have been developed to predict either the sites of metabolism or the metabolites of small molecules, such as FAME 2, MetaPrint2D, Meteor Nexus, ADMET Predictor, and enviPath. These tools usually focus on specific classes of compounds or reaction sets. With the rise of deep learning, more recent tools have focused on predicting untargeted metabolite structures. These approaches are often combined with other tools, such as NMR spectral prediction [5], to enhance the confidence in metabolite identification in human or environmental samples. However, while such tools usually predict extensive lists of potential metabolites, they often lack information about how the predicted metabolites are formed, limiting their interpretability and biological relevance.

BioTransformer was developed as an online tool to help researchers predict metabolites of small molecules with reaction and enzyme information. In BioTransformer 4.0, we implemented a new abiotic module, added over 130 new reactions, and developed a validation module to retain biologically valid metabolites by comparing them with known human metabolites.

## 2. New Features

### 2.1 Abiotic metabolism prediction

We use six different enzyme classes to represent the enzyme that catalyze the reaction: AbioticHydrolysisReactionClass, AbioticReductionReaction Class, SpontaneousReactionClass, DirectPhotolysisReactionClass, Ozonation Class and ChlorinationClass. The reactions rules are collected mainly by converting the reaction library of Chemical Transformation Simulator (CTS) [10]to SMIRKS strings [4] manually, along with some others that are collected from literature reviews. The resulting abiotic reaction database comprises 249 reactions, categorized into four different classes of reactions: 156 photochemistry reactions, 27 chlorination reactions, 31 ozonation reactions and 35 other non-specified reactions.

### 2.2 Prior Biotransformation(See Figure 2)

As shown in Figure 2, when the human body ingests large molecules, such as tannins and starch, it first breaks them down into smaller components through metabolism, which can then be metabolized at various biological locations [1]. To simulate this process, a prior biotransformation module has been developed, which applies a specific set of reaction rules to iteratively decompose the larger compounds into smaller molecules. This prior biotransformation serves as an initial step in all human and gut microbial metabolism modules. The resulting products from this step are then used as substrates for the corresponding downstream modules.

**Figure 1:**
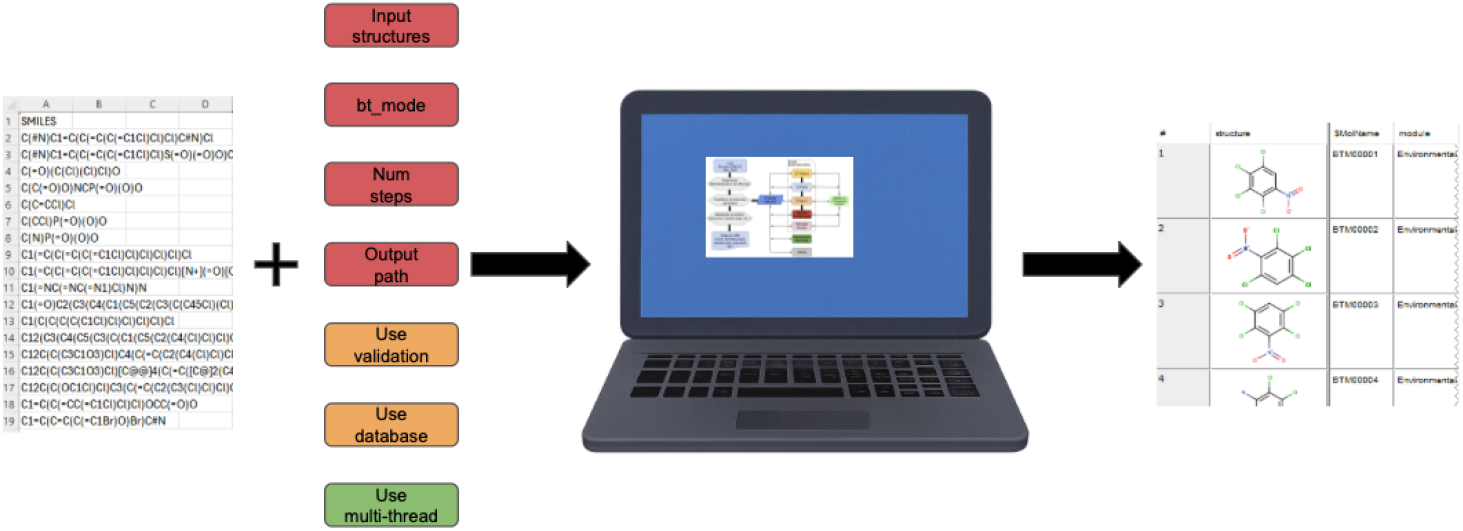
BioTransformer 4.0.

**Figure 2:**
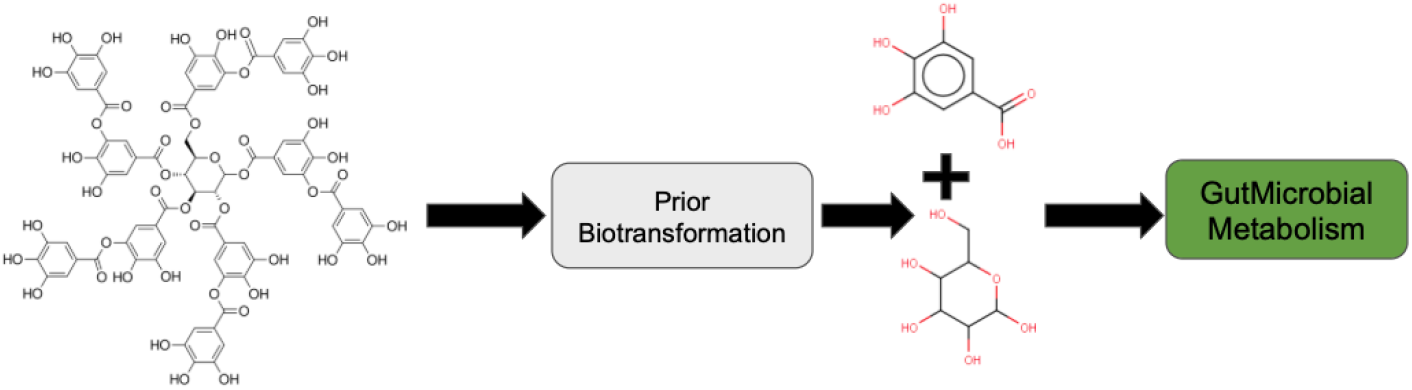
Demonstration of prior biotransformation.

### 2.3 Retrieve metabolites from database

#### 2.3.1 Rhea database in HGut metabolism

Rhea [2] is a reaction database that contains 17783 expert-curated biological reactions that are independent of their biological locations. To enrich the reactions for human gut microbial metabolism, we include 45 reaction rules and 339 substrate metabolite pairs in the [Rhea database](https://www.rhea-db.org/).

#### 2.3.2 Endogenous Handling

Since endogenous compounds and their metabolic fates for human metabolism are well studied, and many participate in numerous metabolism pathways and be transformed to various metabolites. To save computational resource and produce more accurate products for endogenous compounds, we extracted an endogenous database from Pathbank to (1) determine if a compound is endogenous based on the first 14 characters of its InChIKey and (2) return its metabolites by exploring the substrate metabolite pairs in the database. Users can choose to present endogenous metabolites in the saved results or not.

### 2.4 Metabolites Validation

(See figure 3) (See figure 3) In previous versions of BioTransformer, the number of predicted metabolites increased exponentially during multi-step human metabolism prediction, many of which were never observed and are unlikely to be true metabolites. This resulted in computational inefficiencies and made it difficult for users to interpret and validate the predicted results. The metabolites validation module to restrict the metabolites during the prediction process when it’s on. It determines if a metabolite is valid by comparing the structures of itself and its reactive fragment against human metabolites in the validationdb database. The validationdb database stores the first 14 characters of InChIKey, the SMILES string and the pubchem fingerprints of 100059 detected and quan tified, detected but not quantified and expected but not quantified metabolites from HMDB [11] To determine if a metabolite is valid, the program will (1) compute the pubchem fingerprints [8] of the metabolite, (2) extract all candidate structures whose pubchem fingerprint is a superset of metabolite’s from validationdb, (3) extract the reactive fragments from the metabolite by comparing its structure with its substrate; (4) compare if the structures of reactive fragment or the metabolite is a substructure of any candidate structure and return valid if it is or invalid otherwise. By enabling the metabolites validation module, only the valid metabolites are used as substrates for next iteration during multi step metabolism, and the final number of metabolites is reduced to 1/5 to 1/8 in a 6 step metabolism prediction.

**Figure 3:**
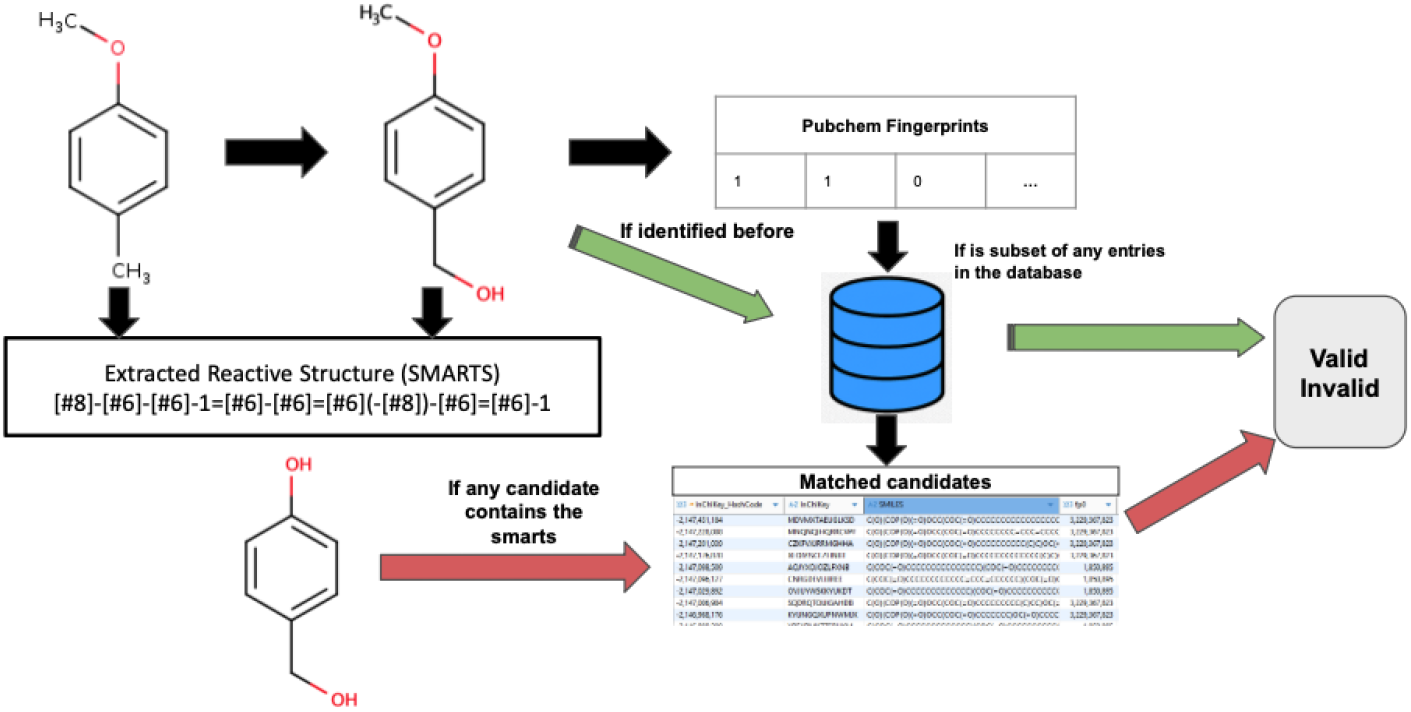
Metabolites Validation.

### 2.5 Metabolic Pathway tool

The metabolic pathway tool builds the metabolic pathways from multi-step predicted results. There are three modes in the metabolic pathway tool, which are substrate mode, product mode and pair mode. 1. The substrate mode takes the structure of a compound in the predicted results as input, and retrieves all metabolic pathways originating from the queried compound. 2. Similarly, the product mode will return all path ways that terminate at the queried compound. 3. For more targeted analysis, the pair model take both a starting substrate and an end product in the results predicted by BioTransformer 4.0 e, the metabolic pathway tool will firstly find the step where the end product is predicted, then tracing back through intermediate the substrate-metabolite pairs iteratively till it reaches the starting substrate. Each metabolic pathway will be returned as a single .csv or .sdf file.

## 3 Results

In this study, we evaluated BioTransformer4.0’s performance by running the six step all-human metabolism prediction on the DrugBank dataset of 2,457 known biotransformations, and the PhytoHub dataset of 633 known biotransformations. The tool achieved recall values of 87.2% and 91.6% for the DrugBank [7] and PhytoHub [3] datasets respectively. The number of predicted metabolites is also reduced from 283 to 52 metabolites per substrate by applying the new developed ontology validator. See figure 4. for an example of the prediction.

**Figure 4:**
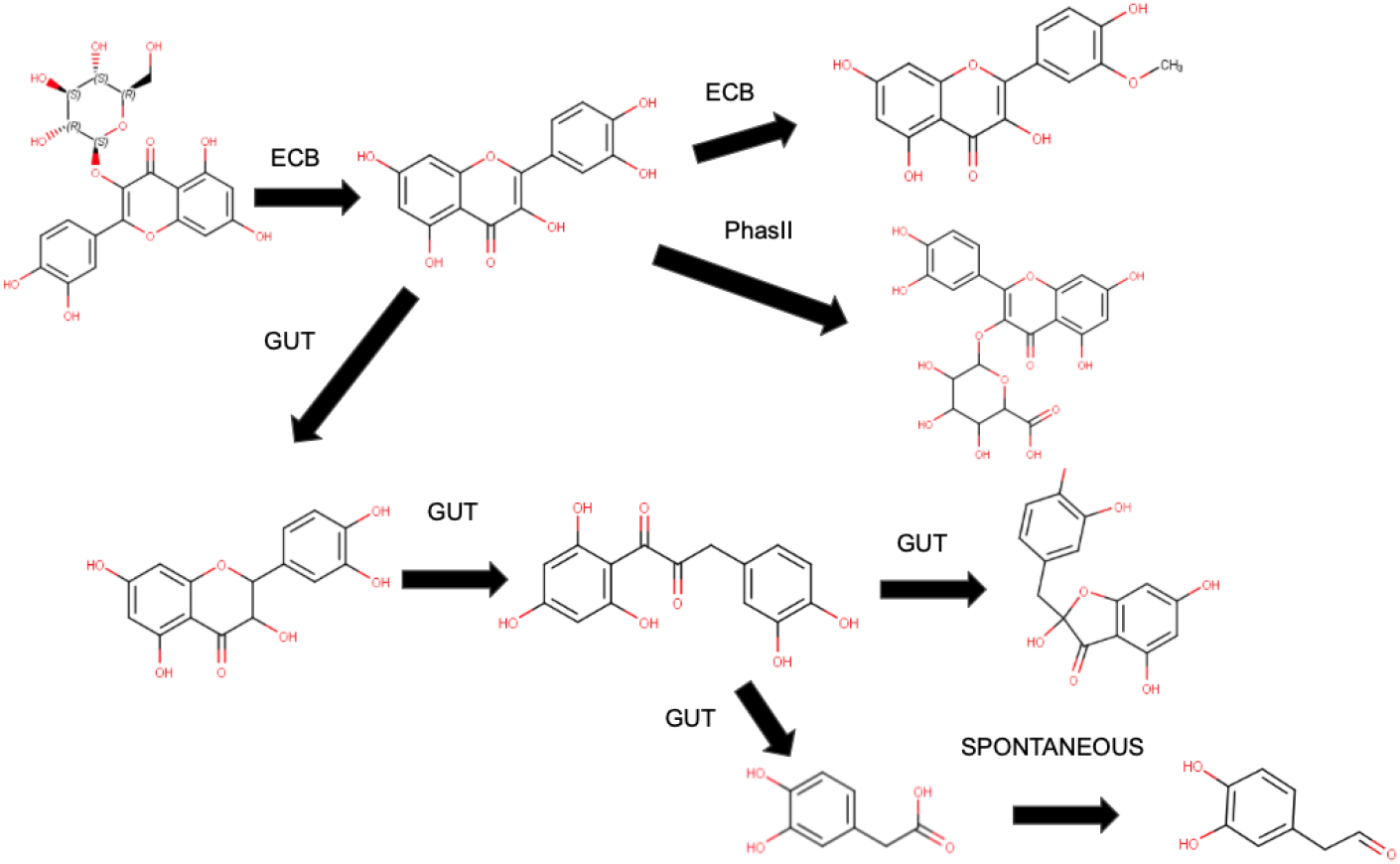
Example of predicted metabolites for isoquercetin.

## 4 Conclusion

BioTransformer 4.0 is a freely available tool that integrates both knowledge based and machine learning approaches to provide more accurate and efficient predictions in small molecule metabolism. The addition of over 130 new reaction rules, along with the inclusion of an abiotic metabolism module, expands the tool’s capabilities to simulate a broader range of biotransformations. Furthermore, the newly developed metabolites validation module has dramatically improved the efficiency and accuracy of predictions. The number of validated predicted metabolites is reduced to about 20% if the raw metabolites, which can help users to review and interpret the results much more efficiently. With its robust performance on both the DrugBank and PhytoHub datasets, achieving recall values of 87.2% and 91.6%, BioTransformer 4.0 stands out as a powerful tool for a wide array of applications, including drug discovery, environmental toxicology, and metabolomics. The customizable multi-step prediction feature, combined with the metabolic pathway tool, enables users to trace complex metabolic pathways. As more metabolites and reactions are identified, and as advancements in deep learning algorithms continue to emerge, we aim to expand BioTransformer by incorporating deep learning techniques and reducing reliance on manually curated reactions. The tool is freely available, and we encourage the scientific community to leverage its capabilities for deeper insights into metabolic processes.

## 5 Disclaimer

We are still finalizing the BioTransformer 4.0. The data and results presented in this preprint are subject to change. Readers are advised to exercise caution when utilizing or interpreting the findings and to await the finalized version for comprehensive validation and updates. The current version may not reflect the final accuracy or completeness of the software, and we recommend that the results be used with caution until further revisions are made.

## References

[1] Bruce Alberts, Alexander Johnson, Julian Lewis, Martin Raff, Keith Roberts, and Peter Walter, editors. Molecular Biology of the Cell. Garland Science, 2002. Available from: NCBI Bookshelf.

[2] Parit Bansal, Anne Morgat, Kristian B Axelsen, Venkatesh Muthukrishnan, Elisabeth Coudert, Lucila Aimo, Nevila Hyka-Nouspikel, Elisabeth Gasteiger, Arnaud Kerhornou, Teresa Batista Neto, et al. Rhea, the reaction knowledgebase in 2022. Nucleic acids research, 50(D1):D693–D700, 2022.

[3] Andreia Bento da Silva, Franck Giacomoni, Balthazar Pavot, Yoann Fillatre, Joseph Rothwell, Bergona Bartolomé Sualdea Catherine Veyrat, Rocio Garcia-Villalba, Cécile Gladine, Rachel Kopec, et al. Phytohub v1. 4: A new release for the online database dedicated to food phytochemicals and their human metabolites. In The 1. International Conference on Food Bioactives & Health, page np, 2016.

[4] Daylight Chemical Information Systems, Inc. SMIRKS: A reaction transform language. https://daylight.com/dayhtml/doc/theory/theory.smirks.html. Accessed: 2025-07-28.

[5] DeepMet. Deepmet. https://deepmet.org, 2024. Accessed: 2025-07-28.

[6] Ricardo Franco-Duarte, Lucia Čerńaková, Snehal Kadam, and et al. Advances in chemical and biological methods to identify microorganisms— from past to present. Microorganisms, 7(5):130, 2019.

[7] Craig Knox, Mike Wilson, Christen M Klinger, Mark Franklin, Eponine Oler, Alex Wilson, Allison Pon, Jordan Cox, Na Eun Chin, Seth A Strawbridge, et al. Drugbank 6.0: the drugbank knowledgebase for 2024. Nucleic acids research, 52(D1):D1265–D1275, 2024.

[8] PubChem Project Team. Pubchem substructure fingerprint specification. https://ftp.ncbi.nlm.nih.gov/pubchem/specifications/pubchem_fingerprints.pdf, 2021. Accessed: 2025-07-28.

[9] Shi Qiu, Ying Cai, Hong Yao, Chunsheng Lin, Yiqiang Xie, Songqi Tang, and Aihua Zhang. Small molecule metabolites: discovery of biomarkers and therapeutic targets. Signal Transduction and Targeted Therapy, 8(1):132, 2023.

[10] U.S. Environmental Protection Agency. Chemical transformation simulator (cts). https://qed.epa.gov/cts/. Accessed: 2025-07-28.

[11] David S Wishart, AnChi Guo, Eponine Oler, Fei Wang, Afia Anjum, Harrison Peters, Raynard Dizon, Zinat Sayeeda, Siyang Tian, Brian L Lee, et al. Hmdb 5.0: the human metabolome database for 2022. Nucleic acids research, 50(D1):D622–D631, 2022.

